# Pharmacokinetics of caffeine: A systematic analysis of reported data for application in metabolic phenotyping and liver function testing

**DOI:** 10.1101/2021.07.12.452094

**Authors:** Jan Grzegorzewski, Florian Bartsch, Adrian Köller, Matthias König

**Author notes:** Correspondence: Matthias König.

## Abstract

Caffeine is by far the most ubiquitous psychostimulant worldwide found in tea, coffee, cocoa, energy drinks, and many other beverages and food. Caffeine is almost exclusively metabolized in the liver by the cytochrome P-450 enzyme system to the main product paraxanthine and the additional products theobromine and theophylline. Besides its stimulating properties, two important applications of caffeine are metabolic phenotyping of cytochrome P450 1A2 (CYP1A2) and liver function testing. An open challenge in this context is to identify underlying causes of the large inter-individual variability in caffeine pharmacokinetics. Data is urgently needed to understand and quantify confounding factors such as lifestyle (e.g. smoking), the effects of drug-caffeine interactions (e.g. medication metabolized via CYP1A2), and the effect of disease. Here we report the first integrative and systematic analysis of data on caffeine pharmacokinetics from 148 publications and provide a comprehensive high-quality data set on the pharmacokinetics of caffeine, caffeine metabolites, and their metabolic ratios in human adults. The data set is enriched by meta-data on the characteristics of studied patient cohorts and subjects (e.g. age, body weight, smoking status, health status), the applied interventions (e.g. dosing, substance, route of application), measured pharmacokinetic time-courses, and pharmacokinetic parameters (e.g. clearance, half-life, area under the curve). We demonstrate via multiple applications how the data set can be used to solidify existing knowledge and gain new insights relevant for metabolic phenotyping and liver function testing based on caffeine. Specifically, we analyzed (i) the alteration of caffeine pharmacokinetics with smoking and use of oral contraceptives; (ii) drug-drug interactions with caffeine as possible confounding factors of caffeine pharmacokinetics or source of adverse effects; (iii) alteration of caffeine pharmacokinetics in disease; and (iv) the applicability of caffeine as a salivary test substance by comparison of plasma and saliva data. In conclusion, our data set and analyses provide important resources which could enable more accurate caffeine-based metabolic phenotyping and liver function testing.

## 1 INTRODUCTION

Caffeine is commonly found in tea, coffee, cocoa, energy drinks, and many other beverages. It is by far the most ubiquitous psychostimulant worldwide (Gilbert, 1984), with 85% of the U.S. population consuming caffeine daily (Mitchell et al., 2014). Among caffeine consumers, the average consumption is more than 200 mg of caffeine per day (Frary et al., 2005). Caffeine is mainly known for its stimulating properties but is also consumed for improved exercise performance and the treatment of various diseases (e.g. apnea in prematurity, hypersomnia). Two important applications of caffeine are liver function testing and metabolic phenotyping of cytochrome P450 1A2 (CYP1A2), N-acetyltransferase 2 (NA2), and xanthine oxidase (XO) (Wang et al., 1985; Jost et al., 1987; Wahlländer et al., 1990; Tripathi et al., 2015).

Caffeine is almost exclusively metabolized in the liver by the cytochrome P450 enzyme system with 3% or less being excreted unchanged in urine (Kot and Daniel, 2008). In humans, N-3 demethylation of caffeine (1,3,7-trimethylxanthine) to paraxanthine (1,7-dimethylxanthine) is the main reaction in the metabolism of caffeine, accounting for around 80-90% of caffeine demethylation. The reaction is exclusively mediated by the activity of the cytochrome P450 isoform 1A2 (CYP1A2) (Hakooz, 2009). The remainder of caffeine is metabolized to around 11% and 4% to the 1-demethylated product theobromine and 7-demethylated product theophylline, respectively (Lelo et al., 1986b; Kalow and Tang, 1993; Miners and Birkett, 1996; Amchin et al., 1999).

Large variation exists in the consumption of caffeine-containing beverages and food between individuals, which can induce CYP1A2 activity. In addition, CYP1A2 activity and protein amount are affected by environmental, genetic, and epigenetic factors (Klein et al., 2010) resulting in large variation between 5-6 fold in humans (Schrenk et al., 1998). These factors lead to a wide range of caffeine plasma concentrations and caffeine pharmacokinetics.

Sex does not significantly influence the CYP1A2 activity (Klein et al., 2010; Yang et al., 2010; Puri et al., 2020). A large heritability of CYP1A2 activity could be shown by a twin study (Rasmussen et al., 2002). However, satisfying genetic or epigenetic markers- on the CYP1A locus on chromosome 15 could not be found by follow-up studies (Jiang et al., 2006; Ghotbi et al., 2007; Gunes and Dahl, 2008; Klein et al., 2010; Yang et al., 2010). Other genes, regulating the expression and function of CYP1A2 and non-genetic factors could explain 42%, 38%, and 33% of CYP1A2 variation at activity, protein, and mRNA level, respectively (Klein et al., 2010). Lifestyle factors (e.g. smoking) and use of oral contraceptives have been shown to influence caffeine pharmacokinetics, as have pregnancy, obesity, alcohol consumption, and the coadministrations of drugs (e.g. fluvoxamine and pipmedic acid). Many diseases reduce the metabolic capabilities of patients. For caffeine which is predominantly metabolized by the liver, various liver diseases result in a strong reduction in caffeine clearance. The most profound reduction is observed in cirrhotic liver disease, correlating with the degree of hepatic impairment (Holstege et al., 1989; Park et al., 2003; Jodynis-Liebert et al., 2004; Tripathi et al., 2015).

Metabolic phenotyping of enzymes by probe drugs is a common method to evaluate the impact of lifestyle, drug-gene and drug-drug interactions, and other factors influencing enzyme activity. Caffeine is an established probe drug for CYP1A2, N-acetyltransferase 2 (NAT2), and xanthine oxidase (XO) metabolic activities (Miners and Birkett, 1996; Fuhr et al., 1996; Faber et al., 2005; Hakooz, 2009). It is rapidly and completely absorbed by the gastrointestinal tract, distributed throughout the total body water, has low plasma binding, as well as short half-life, negligible first-pass metabolism (Yesair et al., 1984), minimal renal elimination, excellent tolerability, and its biotransformation is virtually confined to the liver (Amchin et al., 1999; Kalow and Tang, 1993; Drozdzik et al., 2018). Caffeine is especially used for CYP1A2 phenotyping which contributes 5–20% to the hepatic P450 pool and is involved in the clearance of about 9% of clinically used drugs (Zanger and Schwab, 2013). The partial or systemic caffeine clearance measured in plasma is considered to be the gold standard for CYP1A2 phenotyping (Fuhr et al., 1996) since 95% of the systemic clearance of caffeine is estimated to be due to hepatic CYP1A2 (Amchin et al., 1999). Measurements in serum, saliva, and urine are extensively studied as well. In urine, sampling at multiple time points and precise timing are inherently difficult. Thus, clearance rates are typically calculated only from plasma, saliva, and serum samples. Measurements in saliva are not invasive and show good correlation with measurements in plasma (Callahan et al., 1982; Wahlländer et al., 1990). The metabolic ratio (MR) between various metabolites of caffeine is an established alternative measure for CYP1A2 enzyme activity (Hakooz, 2009). Analogously, the MR is measured in any of the above mentioned tissues though typically only at a single time point after drug administration. The MR of various metabolites at 4 hours after dosing in plasma, saliva, and urine correlate well with the apparent caffeine clearance, 0.84, 0.82, 0.61, respectively (Carrillo et al., 2000). The MRs measured in plasma and urine have been historically most popular. However, measurements in saliva are routinely applied, especially in epidemiological studies (Tantcheva-Poór et al., 1999; Kukongviriyapan et al., 2004; Tripathi et al., 2015; Chia et al., 2016; Urry et al., 2016; Puri et al., 2020).

Despite the great potential of caffeine as a test substance for liver function tests and CYP1A2 based phenotyping, so far caffeine testing has not found widespread clinical adoption. For liver function tests, a major limiting factor is the large inter-individual variability. Data is urgently needed to understand and quantify confounding factors of caffeine pharmacokinetics such as lifestyle (e.g. smoking) and the effects of drug-drug interactions (e.g. drugs metabolized via CYP1A2) or how disease alters caffeine elimination. Based on such data, more accurate liver function tests and CYP1A2 phenotyping protocols could be established. Differences in clinical protocols (e.g. dosing amount, sampling tissue, and timing) have not been systematically analyzed in the literature. In addition, data on competing substances in the context of dynamical liver function tests (e.g. metacethin used in the LiMAx test (Rubin et al., 2017)) and CYP1A2 phenotyping is not accessible but absolutely imperative for a quantitative evaluation of these methods.

Caffeine pharmacokinetics have been investigated in a multitude of clinical trials, each with its own focus and research question. These studies have been reviewed in a broad scope, most recently in (Arnaud, 2011; Nehlig, 2018). Despite caffeine pharmacokinetics being highly studied in literature, no integrated pharmacokinetics data set exists so far and no systematic analysis of the reported data has been performed. The objective of this work was to fill this gap by providing the first comprehensive high-quality data set of reported data on caffeine pharmacokinetics and demonstrate its value via multiple example applications relevant for metabolic phenotyping and liver function testing based on caffeine.

## 2 MATERIAL AND METHODS

### 2.1 Data curation

Publications containing caffeine pharmacokinetics data were searched on PubMed and Google Scholar using combinations of keywords related to caffeine and pharmacokinetics. Most publications were retrieved by combining the keyword caffeine with one of the following keywords: pharmacokinetics, pk, plasma concentration, serum concentration, saliva, time profile, time-course, concentration-time profile, clearance, half-life, elimination rate, or area under the curve (AUC). Based on this initial corpus, additional publications were added by following references and citations. Pharmacokinetics data was curated manually as part of the pharmacokinetics database PK-DB (https://pk-db.com/) (Grzegorzewski et al., 2020) using established workflows. The pharmacokinetics data was stored in combination with relevant metadata on groups, individuals, interventions, and outputs. PK-DB provided support in the curation process with strong validation rules, unit normalization, and automatic calculation of pharmacokinetic parameters from time-courses. As part of the curation process and the presented analyses, pharmacokinetic parameters and other commonly reported measurements were integrated from multiple studies. The meta-analyses and data integration allowed to identify and correct/remove outlier data which were mostly due to either curation errors or incorrect reporting. Pharmacokinetic parameters calculated from time-courses are included in the analyses. For more details see (Grzegorzewski et al., 2020).

### 2.2 Data processing and filtering

Certain data processing and data filtering methods were applied in all analyses. Data included in the analyses was either measured in plasma, serum, or saliva (urinary measurements were excluded). The data was neither filtered based on application route (e.g. oral, intravenous, intramuscular) nor application form (e.g. tablet, capsule, solution) of caffeine. Male subjects were assumed not to take any oral contraceptives. Data outliers from four studies (Stille et al., 1987; Harder et al., 1988, 1989; Balogh et al., 1992) were excluded from all analyses. All four studies probably originate from the same clinical investigation which was detected as outliers during the data curation process.

In Fig. 2A, B and Fig. 3A-D, the data is displayed in a similar manner. For collectively reported subjects, the group size and standard error is displayed as the marker size and error-bar, respectively. In the legend, (I), (G), and (TI) stand for individual participant data, the number of groups, and the total number of subjects, respectively. Data points are depicted as circles if reported equivalently in the source, as squares if calculated from concentration-time profiles and as triangles if inferred from corresponding pharmacokinetic data and body weights of the subjects. Typically, dosing is reported in mass units, AUC in mass per volume units, clearance in volume per time units, and half-life in time units. Occasionally, dosage, AUC, and clearance are reported in body weight units. In case of reported subject weight, the data is harmonized to similar units by multiplying with the reported weights.

**Figure 1.**
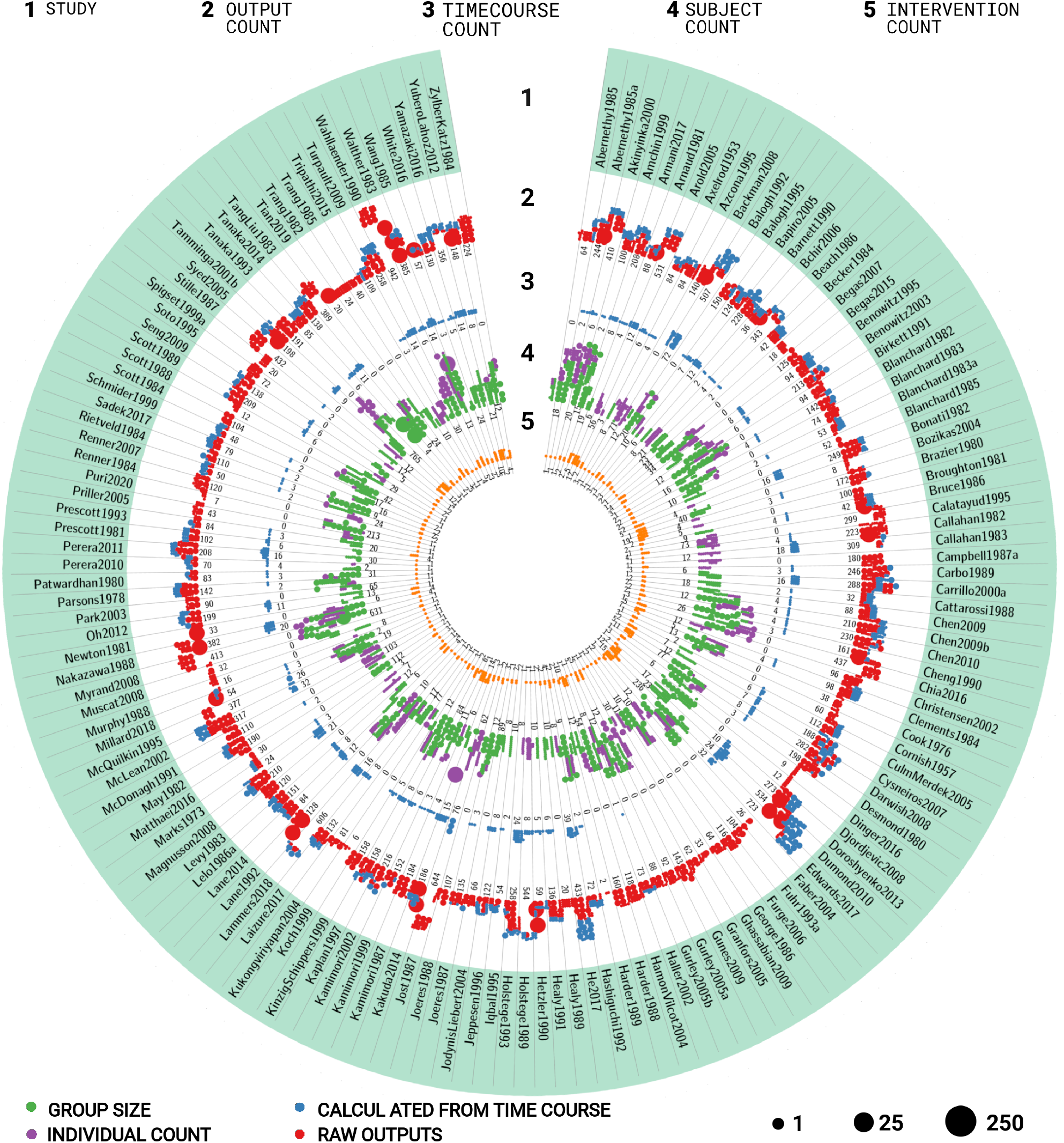
Overview of studies in the caffeine pharmacokinetics data set. The data set consists of 148 studies containing 1311 groups, 12106 individuals, 1650 interventions, 97026 outputs, and 4224 time-courses. The circular plot is structured in stripes and rings. Each stripe represents a different study, each ring the amount of different data types for the respective study. The dots represent the respective amount of data with the dot size corresponding to the number of entries per dot. The rings contain the following information for the respective study (1) name of the study; (2) number of outputs (pharmacokinetics parameters and other measurements). Red dots represent reported data, blue dots data calculated from time-courses reported in the study; (3) number of time-courses; (4) number of participants. Purple dots represent participants with individual data, green dots represent collectively reported participants; (5) number of interventions applied to the participants in the study. For additional information see Tab. 1.

**Figure 2.**
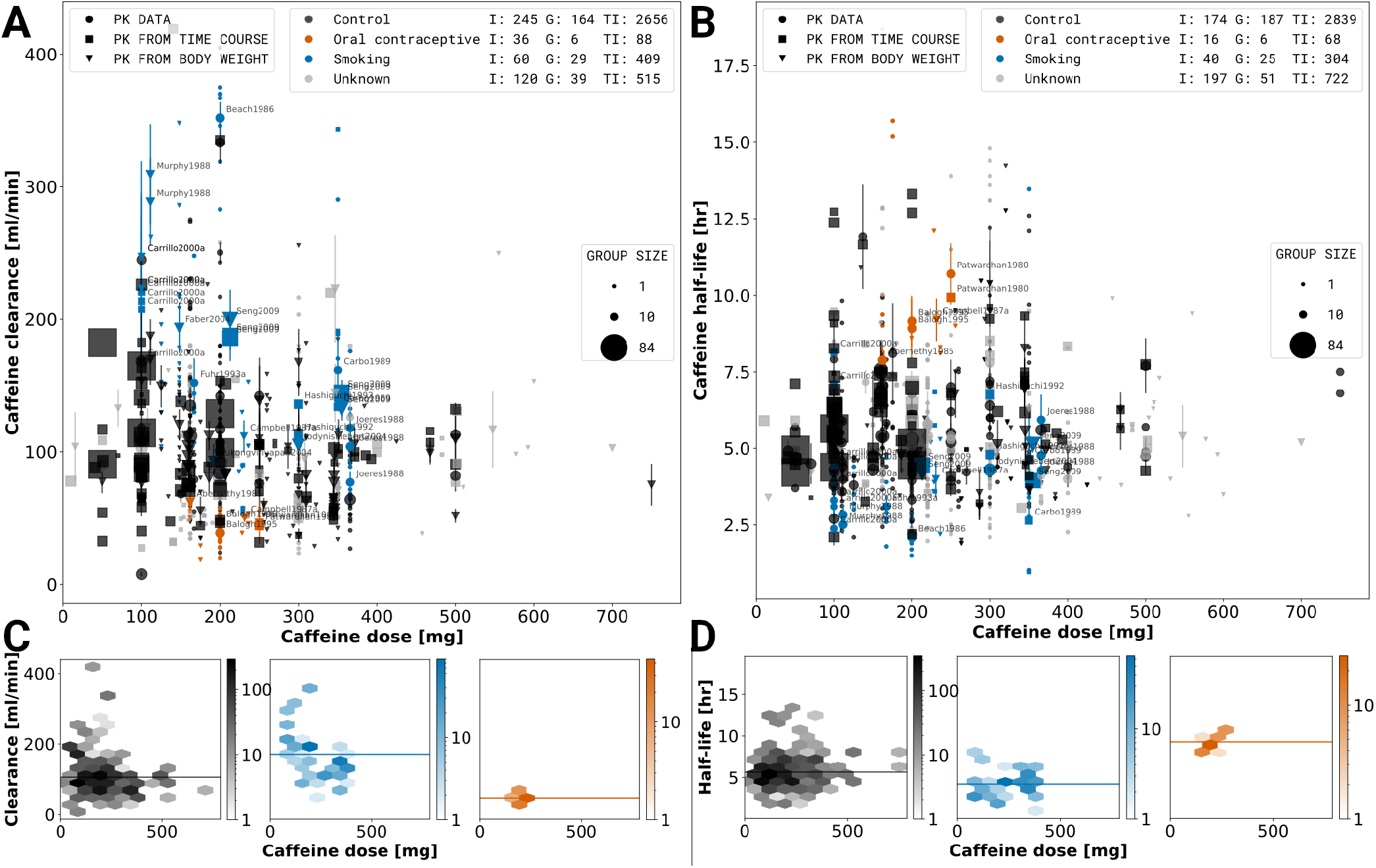
Dose-dependent effect of smoking and oral contraceptive use on caffeine pharmacokinetics. A stratified meta-analysis of caffeine clearance (A) and half-life (B) depending on reported smoking and oral contraceptive use was performed. Black: Control subjects are non-smokers and not taking oral contraceptives; Orange: Oral contraceptive users independent of smoking status (smokers and non-smokers). Blue: Smoking are smokers not consuming oral contraceptives. Grey: Unknown data correspond to subjects with unreported smoking and oral contraceptive status. Marker shape, and size describe the datatype and group size, respectively. Data representing smokers or oral contraceptive consumers is labeled by the respective study name. The hexagonal bin plots in the lower panel (C,D) correspond to the subset of data for the control, smoking, and oral contraceptive consuming subjects. The color intensity of each bin represents the number of subjects falling in a given hexagonal bin area. Data selection criteria and visualization are described in the methods section.

**Figure 3.**
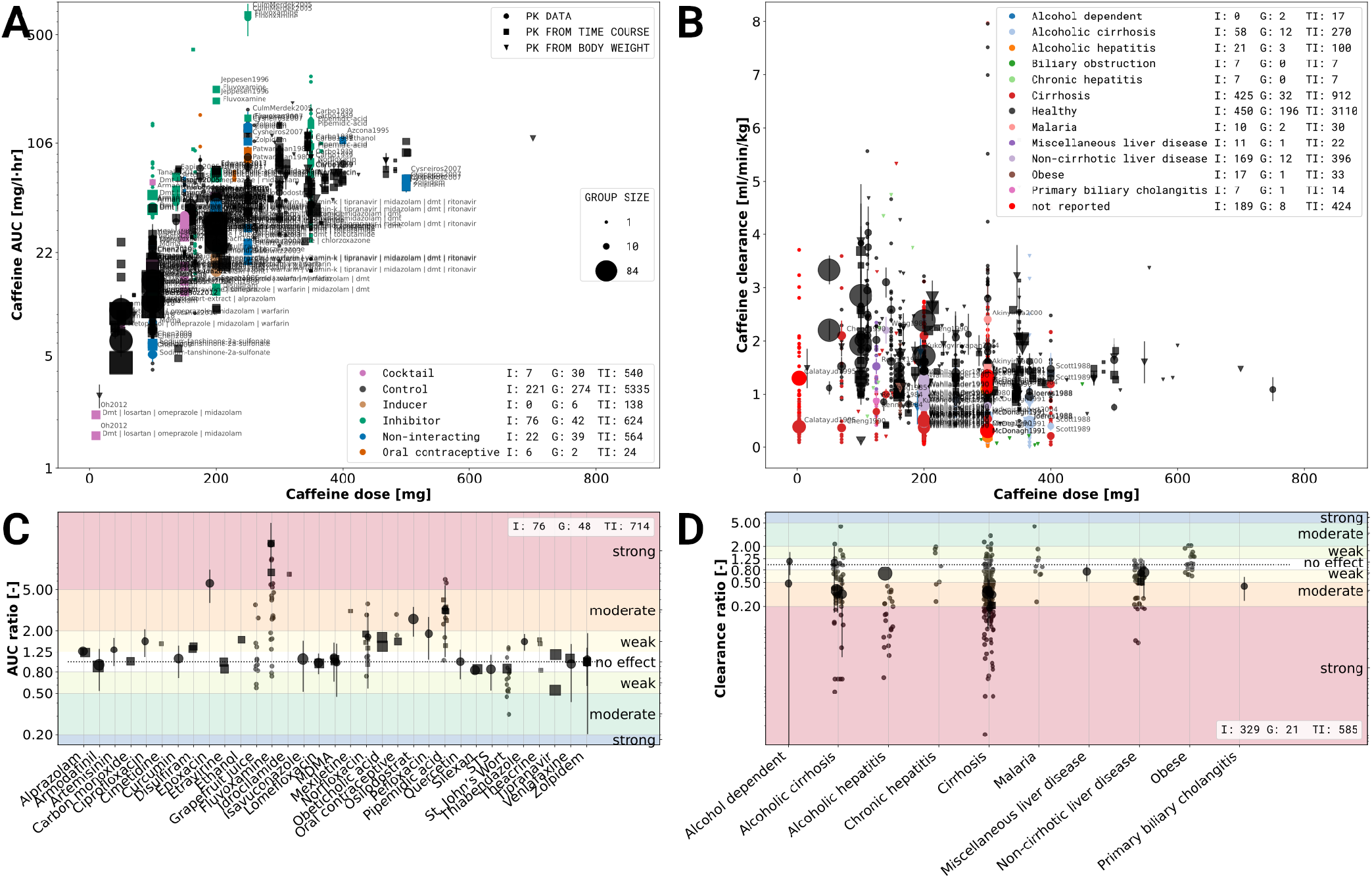
Effects of caffeine-drug and caffeine-disease interactions. A) Caffeine-drug interactions based on the caffeine area under the concentration curve (AUC). Data is stratified based on co-administration of drugs with caffeine. Violet: caffeine administrated as part of a drug cocktail. Common co-administrations are dextrometorphan, metoprolol, midazolam, omeprazole, and warfarin.; Black: single caffeine administration (no co-administration); Brown: co-administration with an inducing effect on the elimination of caffeine; Green: co-administration with an inhibiting effect on the elimination of caffeine; Blue: co-administrations with no effect on the pharmacokinetics of caffeine; Orange co-administration of oral contraceptives. B) Caffeine-disease interactions based on caffeine clearance. Data was stratified based on the health status and reported diseases, with black data points corresponding to healthy subjects. C) Effect sizes of caffeine-drug interaction for studies with a controlled study design, mostly randomized control trials (RCT). The effect size is based on the log AUC ratio between caffeine application alone and caffeine with co-administration of the respective drug. The drugs were characterized as having either a strong, moderate, weak or no effect. Strong, moderate and weak inhibitors increase the AUC ≥ 5-fold, ≥ 2 to < 5-fold, ≥ 1.25 to < 2-fold, respectively. D) Effect size of caffeine-disease interactions for studies with a controlled study design, mostly case-controlled studies. The effect size is based on the log clearance ratio between subjects with and without a specific condition/disease. The diseases were characterized as having either a strong, moderate, weak or no effect. Strong, moderate and weak effect decreased the clearance by ≥ 80 percent, ≥ 50 to < 80 percent and ≥ 20 to < 50 percent, respectively. Data selection criteria and visualization are described in the methods section.

In Fig. 2, the depicted subjects were healthy. Substances with negligible caffeine-drug interactions were determined by an effect size analysis in Sec. 3.4. All other co-administrations and investigations containing multiple caffeine dosages were excluded.

In subplot Fig. 3A, included subjects were healthy. The area under the caffeine concentration curves measured at least up to 12 hours after a single application of caffeine and AUC extrapolated to infinity were included. Multiple subsequent caffeine dosages were excluded, other administrations and co-administrations included. In subplot Fig. 3C, healthy and non-healthy subjects were included. Single applications of caffeine or caffeine administrated as a cocktail with negligible drug-drug interactions were included. All other co-administrations and investigations containing multiple caffeine dosages were excluded.

In subplots Fig. 4A-C, no data was excluded. In subplot Fig. 4D, included subjects were healthy, non-smoking, non-pregnant, and non-oral contraceptive consumers. Interventions with caffeine administrated as a cocktail with negligible caffeine-drug interactions were included. All other co-administrations and investigations containing multiple caffeine dosages were excluded.

**Figure 4.**
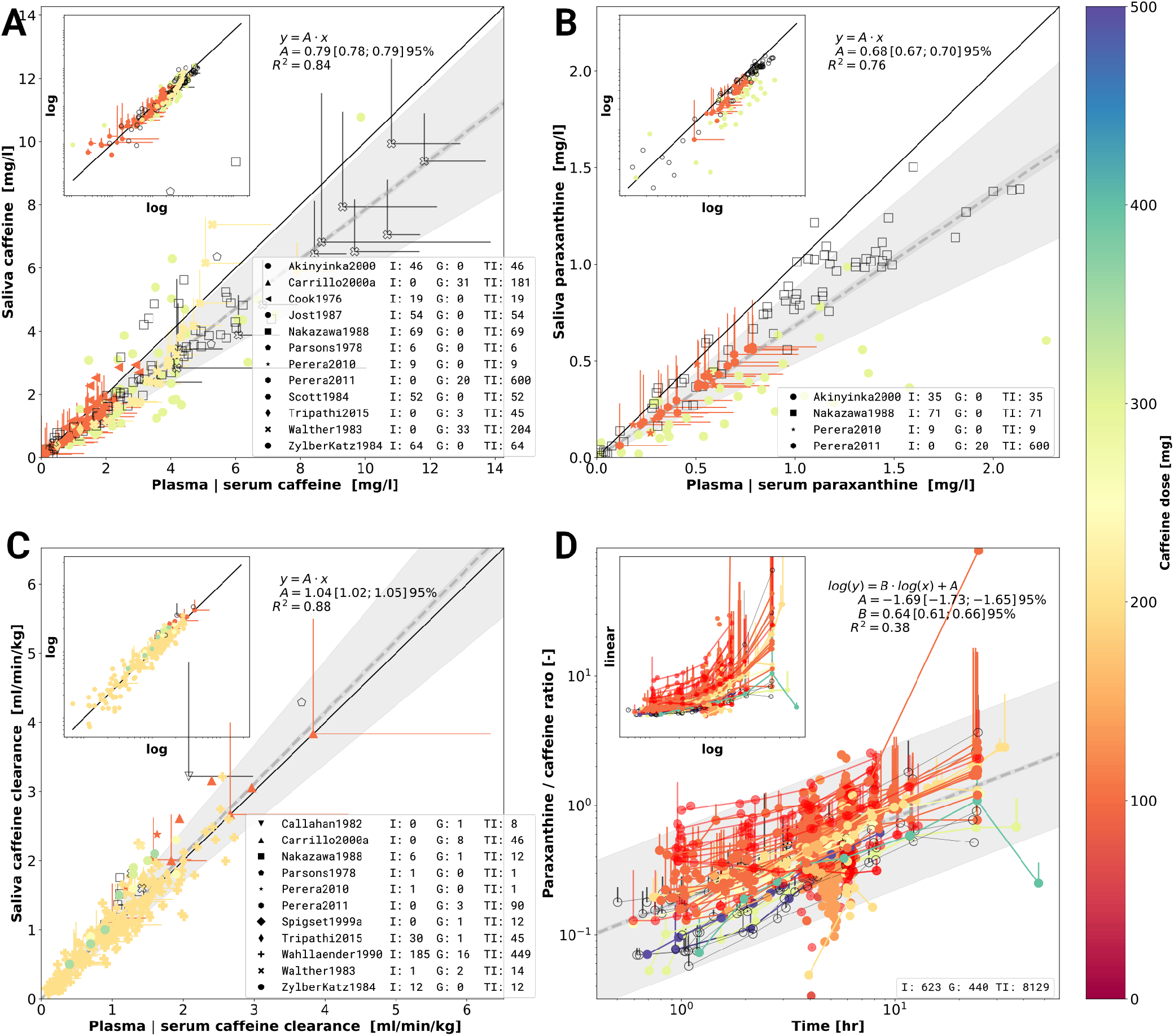
Meta-analysis of caffeine and paraxanthine concentrations in plasma, serum and saliva. A) Caffeine concentrations in saliva versus caffeine concentrations in plasma or serum. Individual data points come from a single investigation taken at identical times after caffeine dosing. Marker shape encodes the different study. Markers are color-coded by caffeine dose in mg. Empty markers correspond to data in which dosage was reported per body weight but no information on the subject weight was available. B) Paraxanthine concentrations in saliva versus paraxanthine concentrations in plasma or serum analogue to A). C) Caffeine clearance calculated from caffeine concentrations in saliva versus plasma or serum clearance. The panels A, B and C are in linear scale with a log-log inlet showing the same data. The dashed line in A, B and C represents a linear regression (*y* = *A · x*) with wide shaded area being 95% confidence interval of the sample variability and narrow shaded area the 95% confidence interval of the fitted mean of the scaling factor *A*. D) Time dependency of the metabolic ratio paraxanthine/caffeine. Metabolic ratios are measured in plasma, serum, or saliva. Data points belonging to a single time course from a study are connected via a line. The dashed line corresponds to the linear regression *log*(*y*) = *B · log*(*t*) + *A*. Jitter was applied on the time axes for better visibility of overlapping points. Data selection criteria and visualization are described in the methods section.

## 3 RESULTS

### 3.1 Caffeine pharmacokinetics data set

Within this work, the first comprehensive open pharmacokinetics data set on caffeine was established. We demonstrate the value of the data set by its application to multiple research questions relevant for metabolic phenotyping and liver function testing: (i) the effect of smoking and oral contraceptive use on caffeine elimination (Sec. 3.3); (ii) the interaction of caffeine with other drugs (Sec. 3.4), (iii) alteration of caffeine pharmacokinetics in disease (Sec. 3.5) and (iv) the applicability of caffeine as a salivary test substance by comparison of plasma and saliva data (Sec. 3.6). The data set integrates data from 148 publications (Fig. 1 and Tab. 1), with most of the publications corresponding to a distinct clinical trial. The focus of data curation was on pharmacokinetics data of caffeine, caffeine metabolites, and caffeine metabolic ratios in human adults. Importantly, the data set is enriched with meta-data on (i) the characteristics of studied patient cohorts and subjects (e.g. age, body weight, smoking status, health status, fasting); (ii) the applied interventions (e.g. dosing, substance, route of application); (iii) measured pharmacokinetic time-courses; and (iv) pharmacokinetic parameters (e.g. clearance, half-life, area under the curve). In summary, data from 1311 groups and 12106 individuals is reported under 1650 interventions resulting in 97026 pharmacokinetic outputs and 4224 time-courses. The data set is available via the pharmacokinetics database PK-DB (https://pk-db.com/) with a detailed description of the data structure provided in (Grzegorzewski et al., 2020). In the following, we summarize the quality of reporting and provide example applications of the data set.

**Table 1.**
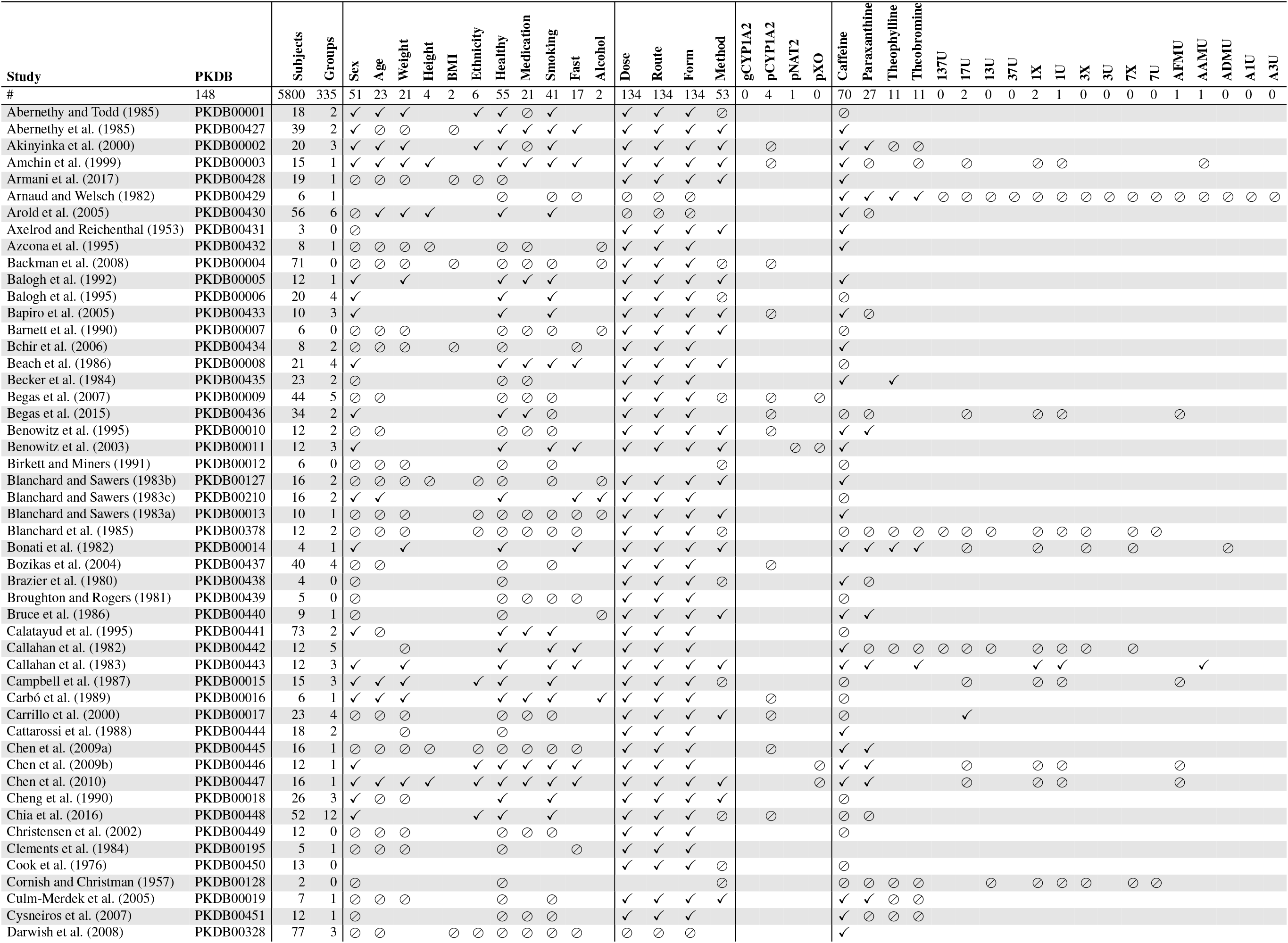
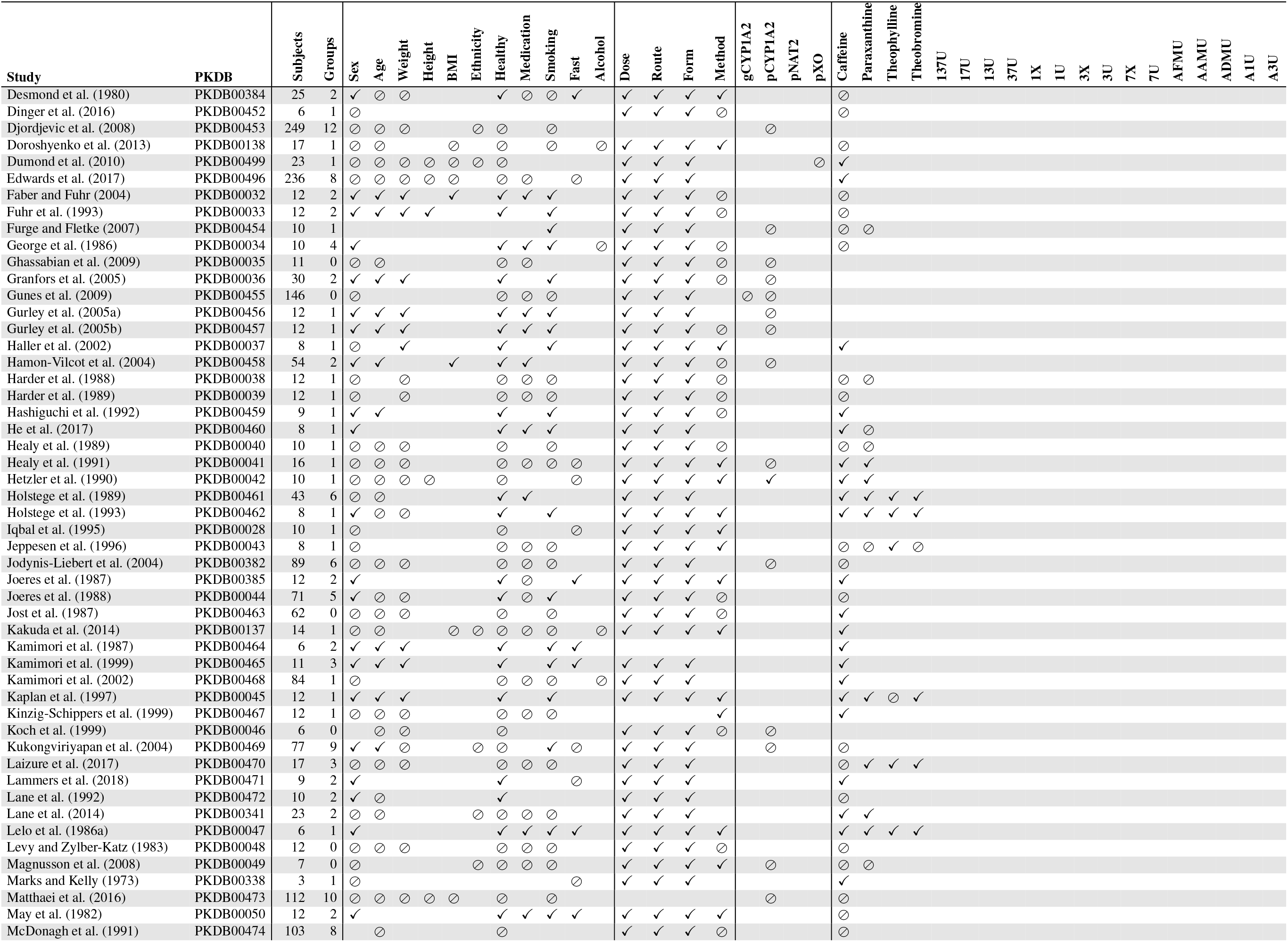
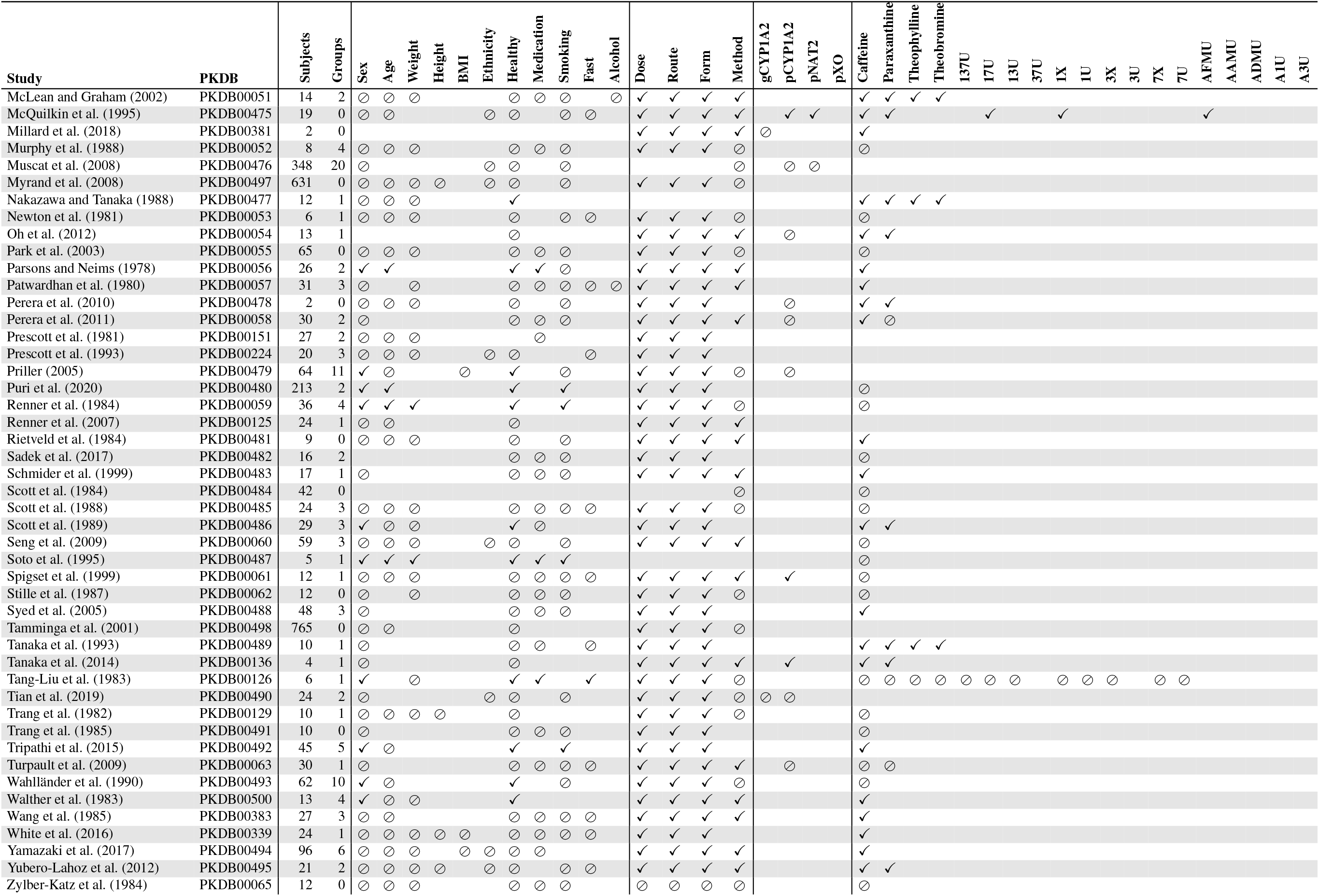
Overview of curated studies. For each study the table shows which information was reported. Either, information was reported (✓), partially reported (⊘) or not reported at all (whitespace).

### 3.2 Reporting of pharmacokinetics data

The study design as well as quality and details of reporting of results were very heterogeneous between studies. Major differences exist in the study design, number of study participants, and number of reported time-courses. Many studies report some individual participants data (84/148) but only a minority of the studies report individual participant data for all study participants (6/84) (Axelrod and Reichenthal, 1953; Brazier et al., 1980; McQuilkin et al., 1995; Millard et al., 2018; Perera et al., 2010; Rietveld et al., 1984). Many studies report only aggregated data on group level (64/148). In most studies, the application of a single dose of caffeine was studied (126/148). In the case of multiple interventions (55/148), mostly one additional substance was co-administrated (38/148). The main categories of studies were either (i) case-control studies which compare caffeine pharmacokinetics in two groups (e.g. healthy vs. disease) (62/148), (ii) crossover studies on caffeine-drug interactions (comparing caffeine alone vs. caffeine and additional substance) (38/148) (iii) studies on metabolic phenotypes (including drug cocktails) (37/148); or (iv) methodological studies (e.g. establishing mass spectroscopy protocol for quantification or new site of sampling).

Intervention protocols, i.e. the applied substances, form, dose, and timing of application was typically reported in good detail. In crossover studies, the difference between treatments was generally reported in good detail. For the dosing with caffeine, the amount (140/148), route (e.g. oral, intravenous) (145/148), form (tablet, capsule, solution) (142/148), and the substance (148/148) are typically reported. Co-administrations of medication and other substances are often mentioned qualitatively (27/55), skipping either the amount, route, form, or exact timing of application.

The quantification protocol, i.e. quantification method (e.g. high-performance liquid chromatography, gas-liquid chromatography) (97/148), the site of sampling (e.g. plasma, serum, saliva, urine) (148/148), and the time points when samples were taken (129/148), were mostly reported in good detail. However, in some studies the quantification method and protocol are not mentioned explicitly but only via additional references, which complicates the curation.

The information on subject characteristics was less often reported in sufficient detail with large differences in the quality and quantity of reporting between studies. Any information on sex (137/148), weight (81/148), and age (94/148) was relatively often reported on group or individual level. However, age and weight were rarely reported on an individual level and often not even for all groups. Other anthropometric factors such as height (15/148), body mass index (BMI) (15/148), and ethnicity (26/148) are rarely reported. The genotype of CYP1A2 (gCYP1A2) is almost never reported (3/148), even though there is evidence that genetic variation can play an important role in caffeine metabolism. It is worth noting that low-cost genotyping methods were not available for early studies included in the data set. The phenotype (pCYP1A2, pXO, pNAT2) of enzymes involved in the metabolism of caffeine, i.e., CYP1A2, xanthine oxidase (XO), or N-acetyltransferase type 2 (NAT2) were investigated occasionally (37/148). The information on other factors influencing the pharmacokinetics of caffeine is reported very heterogeneously. The strong influence of smoking (106/148) and the use of oral contraceptives (42/148) on the enzyme activity of CYP1A2 and thereby on the apparent clearance of caffeine is covered relatively well in many publications. Health status and patient diseases are often covered (140/148). However, often categorized in broad and general disease classes, with more specific disease classification lacking. Markers related to cardiovascular health (e.g. blood pressure, cholesterol level, bilirubin level) are basically not reported in the context of caffeine pharmacokinetics. In case of cirrhosis, further information of severity is reported sparsely. Important information on the abstinence of caffeine or methylxanthines and consumption of caffeine or other caffeine-containing beverages is often missing.

Individual-level reporting is essential for subsequent pharmacokinetic modeling, as biological mechanisms responsible for the pharmacokinetics strongly correlate with these factors and large inter-individual variability exists in caffeine pharmacokinetics. Despite the importance of individual subject data, information on individuals is rarely provided.

Most studies report pharmacokinetics outputs on caffeine (127/148). Data on the main product paraxanthine (46/148) and secondary metabolites theobromine (20/148) and theophylline (19/148) are reported sometimes. Additional metabolites such as 137U, 17U, 13U, 37U, 1X, 1U, 3X, 3U, 7X, 7U, AFMU, AAMU, ADMU, A1U, and A3U are seldom reported, mostly as urinary measurements. In summary, we established the largest freely available pharmacokinetics data set of caffeine so far consisting of data from 148 publications and clinical trials. By using a highly standardized data curation pipeline (established as part of PK-DB (Grzegorzewski et al., 2020)) high quality and consistency of the data could be achieved. This enabled us to apply the data set to various research questions relevant for metabolic phenotyping and liver function tests despite the heterogeneity in reporting. In the following, we demonstrate the value of the data set via multiple example applications consisting of (i) stratification of caffeine pharmacokinetics due to smoking and oral contraceptive use (Sec. 3.3), (ii) the interaction of drugs with caffeine (drug-drug interaction; Sec. 3.4); (iii) alteration of caffeine pharmacokinetics in disease (drug-disease interaction, Sec. 3.5; and (iv) metabolic phenotyping and liver function testing (Sec. 3.6).

### 3.3 Smoking and oral contraceptives

In a first analysis, we were interested in the effect of smoking and oral contraceptive use on the pharmacokinetics of caffeine in healthy subjects (see Fig. 2). Both have repeatedly been reported as key exogenous factors affecting caffeine elimination. A main question was how reproducible the effect is and if by integrating data from multiple studies a more consistent picture of the effects can be gained. For the analysis the data set was stratified into smokers, oral contraceptive users and a control group (neither smoking nor using oral contraceptives). Smoking results in increased caffeine clearance (Fig. 2A,C) and decreased half-life of caffeine elimination (Fig. 2B,D) whereas oral contraceptive use has the opposite effect over a wide dose range of caffeine. An important result from our analysis is that a consistent and reproducible effect can be found over more than 50 years of pharmacokinetic research. With exception of a few outlier studies probably from a single clinical trial (see methods) all data was highly consistent. This provides a strong argument for the applied methods and protocols.

Despite the large effect of smoking and oral contraceptive use on the pharmacokinetics of caffeine, the information is only reported for a subset of studies. Smoking as well as oral contraceptive use should be an exclusion criteria for subjects in studies of caffeine pharmacokinetics, due to the possible confounding effects. Importantly, in many groups smokers and non-smokers were mixed without reporting data for smokers and non-smokers separately. Without reporting of data on individuals or subgroups no stratification could be performed, which could strongly affect results if not balanced between groups. In summary, the integrative data analysis showed a consistent strong activating effect of smoking on caffeine elimination and an inhibiting effect of oral contraceptive use on caffeine elimination.

### 3.4 Caffeine-drug interactions

An important question for metabolic phenotyping and liver function testing with caffeine is how the coadministration of other drugs and compounds affects caffeine clearance. Consequently, we studied in the second analysis the reported caffeine-drug interactions in the data set. The impact of drugs was quantified using the change in AUC between a coadministration and a respective control 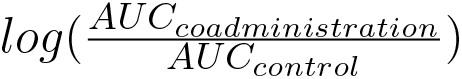 (see Fig. 3A,C).

Only case-controlled studies, mostly cross-over trials with a washout phase were included in the analysis. Corresponding controls were not matched across different studies. Overall coadministration data with AUC difference was available for 33 substances in our data set. In accordance with FDA, EMA and PMDA guidelines (Sudsakorn et al., 2020) we classified substances as inhibitors or inducers of caffeine clearance based on changes in AUC: FDA - Clinical DDI guidance: strong, moderate and weak inhibitors increase the AUC ≥ 5-fold, ≥ 2 to < 5-fold, ≥ 1.25 to < 2-fold, respectively; strong, moderate and weak inducers decrease the AUC by ≥ 80 percent, ≥ 50 to < 80 percent and ≥ 20 to < 50 percent, respectively.

Most substances do not affect the AUC of caffeine, with the exception of fluvoxamine, pipemidic acid and norfloxacin, which inhibit caffeine clearance. Tipranavir was the only substance showing a weak induction of caffeine clearance, but only in steady state dosing (not after a single dose) (Dumond et al., 2010)). Substances administered as a cocktail along side caffeine, aiming to phenotype several enzymes simultaneously did not affect the AUC of caffeine, substantiating the use of caffeine as part of a drug cocktail design (Armani et al., 2017; Dumond et al., 2010; Doroshyenko et al., 2013; Edwards et al., 2017; Kakuda et al., 2014; Lammers et al., 2018; Oh et al., 2012; Tanaka et al., 2014; Turpault et al., 2009). Our analysis shows that only a minority of studied drugs show an interaction with caffeine, confirming its value for phenotyping even under co-administration. As a side note, the protocols for studying caffeine-drug interactions were highly variable, e.g., the applied caffeine dose and the dose of the coadministrated substance varied between studies. Our results suggest that most medications can be safely consumed in combination with caffeine with exception of the antidepressant fluvoxamine, the antibacterial pipemidic acid and the antibiotic norfloxacin in which case caution is warranted.

### 3.5 Caffeine-disease interactions

An important question for using caffeine as a test substance for liver function testing and phenotyping, as well as for drugs metabolized via CYP1A2, is how disease affects the pharmacokinetics and elimination of caffeine. To study this question, we stratified caffeine clearance rates based on the reported disease of subjects and groups in the data set. To quantify the effect of disease the absolute clearance of caffeine (Fig. 3B) and the logarithmic difference to a control group 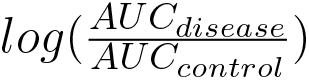 (Fig. 3D) were analyzed. The corresponding controls were not matched across different studies.

None of the reported diseases increased the clearance rate of caffeine. Most of the diseases contained in this data set are diseases of the liver (e.g. alcoholic cirrhosis, primary biliary cholangitis) or are known to affect the liver (e.g. alcohol dependent). Cirrhotic liver disease had moderate to strong effects on the caffeine clearance with large variability in the reported data. Malaria and obesity had no effect on clearance with caffeine. An issue in the study of caffeine-disease interaction is that control group and disease group are different subjects (no cross-over design). In addition diseases were reported very heterogeneously (e.g. either only cirrhosis or with underlying cause such as alcoholic cirrhosis).

### 3.6 Metabolic phenotyping

An important question for metabolic phenotyping and liver function tests with caffeine is how saliva measurements correlate with plasma or serum caffeine measurements, in the following referred to as blood-based measurements. A good correlation would allow simple non-invasive phenotyping using saliva samples. To study this question we analyzed (i) the relationship of blood-based concentrations of caffeine and paraxanthine with their respective saliva concentrations (Fig. 4A,B); and (ii) how the caffeine clearance measured in saliva correlate to blood-based measurements (Fig. 4C).

Systematic errors due to different dosing protocols and different clinical investigation seem to be minimal as the data from multiple studies shows very consistent results. Linear regressions were performed to quantify the relation between saliva and blood-based caffeine and paraxanthine measurements (see Fig. 4A-C). The resulting scaling factors of saliva to blood-based concentration of caffeine and paraxanthine are 0.79 ± 0.01 and 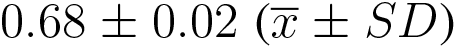, respectively. Pearson correlation coefficients between saliva and blood-based concentrations for caffeine and paraxanthine are 0.84 and 0.76, respectively.

When comparing saliva-based caffeine clearance against blood-based clearance (Fig. 4C) an even stronger correlation of 0.88 with a scaling factor of 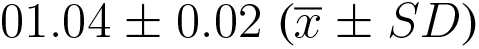 is observed. The integrated clearance data strongly indicates that clearance can either be calculated from saliva or blood-based measurements.

Paraxanthine/caffeine ratios are mainly used for metabolic phenotyping based on caffeine. Whereas most studies use 6 hours to phenotype no clear consensus exists in the literature and metabolic ratios are reported for varying time points after caffeine application. Paraxanthine/caffeine ratios for caffeine administered either as a single dose or in a cocktail to healthy, non-smoking, non-pregnant, and non-oral contraceptive consuming subjects were investigated (Fig. 4D). By applying this strict data filtering, the variability due to either smoking and oral contraceptive use (see Sec. 3.3), caffeine-drug interactions (see Sec. 3.4) or disease (see Sec. 3.5) could be removed from the metabolic ratios.

Early and late time-sampling are least suitable for phenotyping. At these time points concentrations are low, resulting in relatively high random errors and thus low single to noise ratio. In an early stage, the outcome of metabolic ratios are further influenced by the distribution phase of the substance and its absorption kinetics, both affected by form and route of administration. Main results are that the metabolic phenotyping with paraxanthine/caffeine ratios is strongly time dependent with increasing ratios with time; and that a clear caffeine-dose dependency exists in the phenotyping with smaller caffeine doses increasing the metabolic ratio. Our results show the importance of clear standardized protocols for metabolic phenotyping.

In summary, by integrating data from multiple studies we could show a very good correlation between saliva and plasma caffeine concentrations, paraxanthine concentrations and pharmacokinetic parameters calculated from saliva versus plasma concentrations. This pooled data provides a strong argument for caffeine phenotyping based on saliva samples. Furthermore, we analyzed the time dependency of paraxanthine/caffeine ratios often used for phenotyping of CYP1A2. Our time dependent correlation allows to correct caffeine/paraxanthine ratios depending on the time after application and caffeine dose.

## 4 DISCUSSION

Within this work, we performed a systematic data integration and multiple data analyses of reported data on caffeine pharmacokinetics in adults with focus on applications in metabolic phenotyping and liver function testing. To our knowledge, this is the largest data set and analysis of its kind so far. Data integration from multiple sources allows to solidify existing knowledge, increases statistical power, increases generalizability, and creates new insights into the relationship between variables (Thacker, 1988). For instance, by systematically curating group and subject information on smoking status and oral contraceptive use, we could show a reproducible and consistent effect of smoking induction and inhibition via oral contraceptives over many studies and the complete dose regime of applied caffeine. Only a small subset of studies was specifically designed to study these questions (e.g. smoking by (Benowitz et al., 2003; Gunes et al., 2009; Parsons and Neims, 1978; Backman et al., 2008) and oral contraceptives by (Patwardhan et al., 1980; Abernethy and Todd, 1985; Rietveld et al., 1984)). Publicly providing comprehensive pharmacokinetics data in combination with detailed metadata allows to study new aspects of caffeine pharmacokinetics often not even anticipated by the original investigators. One example of such a new aspect is the determination of dose and time-dependency of metabolic phenotyping via paraxanthine/caffeine ratios besides many of the data sets only reporting data for a single time point and dose. Naturally, the advantages of open data are not exclusive to the study of caffeine, but apply to any given substance.

Like all data integration and meta-analysis methods limitations due to sampling bias, biased outcome interpretation and inadequate data may exist (Thacker, 1988). Despite being the most comprehensive analysis so far, we could only present selected aspects mainly driven by the availability of reported data. Our focus in this study was on key factors affecting caffeine pharmacokinetics (smoking and oral contraceptive use), caffeine-drug and caffeine-disease interactions, as well as information relevant for the metabolic phenotyping and liver function testing with caffeine. Other important factors such as the pharmacogenetics of caffeine or urinary metabolic ratios have not been presented. Importantly, the corresponding data has been curated and is readily available in PK-DB, but often very sparse (e.g. in case of genetic variants) or very heterogeneous (e.g. in case of urinary data). Despite many implemented measures to ensure high data quality (e.g. validation rules and checking of studies by multiple curators), we are aware that the created data set may contain mistakes. Please report such instances so that these can be resolved.

The scope of the presented data set is limited to caffeine pharmacokinetics in human adults. Future work will extend our data curation effort towards children and infants and include studies with animal experiment. In addition, despite trying to be as comprehensive as possible additional studies with relevant data exist. Please contact us in such cases so that we can include these additional studies. Contribution of missing data is highly appreciated. Also if you want to contribute a caffeine data set of your own please get in contact.

The large inter-individual variability in caffeine pharmacokinetics is a major limitation for metabolic phenotyping and liver function tests. Our work allowed to evaluate the effect of key factors on caffeine pharmacokinetics such as smoking, oral contraceptive usage or caffeine-drug interactions. An important next step will be the development of methods to quantify and correct for these confounding factors. This could allow to reduce variability in caffeine based testing. A promising tool in this context are physiological-based pharmacokinetics (PBPK) models (Jones and Rowland-Yeo, 2013) using information from the established data set as input for stratification and individualization. We could recently show that such an approach based on a similar data set for indocyanine green (ICG) allowed to account for important factors affecting ICG based liver function tests (Köller et al., 2021).

An important outcome of our analysis are the very good correlations between saliva- and blood-based measurements for caffeine with 0.77 *R*^2^=0.74 in very good agreement with data reported previously in individual studies 0.74 ± 0.1 *R*^2^=0.98 (Akinyinka et al., 2000), 0.79 ± 0.05 *R*^2^=0.96 (Zylber-Katz et al., 1984), 0.74 ± 0.08 (Newton et al., 1981), 0.74 ± 0.08 *R*^2^=0.90 (Jost et al., 1987), 0.71 *R*^2^=0.89 (Scott et al., 1984), 0.73 ± 0.06 (Walther et al., 1983), and for paraxanthine 0.68 *R*^2^=0.76 compared to 0.77 *R*^2^=0.91 (Nakazawa and Tanaka, 1988).

Several studies have shown that blood-derived pharmacokinetic parameters show excellent correlation with saliva-derived parameters (Akinyinka et al., 2000; Newton et al., 1981). We could confirm this observation when systematically analyzing the correlation between saliva- and plasma/serum-derived clearance. Overall we could show that the data from multiple studies are in very good agreement with each other after excluding data with confounding factors such as smoking or oral contraceptive use. These integrated results are a strong argument for saliva based metabolic phenotyping and liver function tests with caffeine, with sampling from saliva being convenient, painless, economical, without the requirement for special devices. Further, they allow simple repeated sampling as often required for pharmacokinetic research (Zylber-Katz et al., 1984).

Importantly, our results are not only applicable to caffeine, but many aspects can be translated to other substances metabolized via CYP1A2, e.g. to the LiMAx liver function tests based on the CYP1A2 substrate methacetin (Rubin et al., 2017). E.g. based on our analysis we would expect smoking and oral contraceptive use to be confounding factors of LiMAx tests, which should be recorded and be accounted for in the evaluation of liver function.

Our systematic curation and analysis of reported caffeine data provided an overview of the current state and limitations of reporting of pharmacokinetic data. In summary, an accepted standard, minimum information guidelines, and standardized meta-data for the reporting of pharmacokinetics data of caffeine are missing. This finding is apparently not only true for the pharmacokinetics of caffeine but rather generally true for reporting of pharmacokinetics research. Major shortcomings in reporting are missing minimum information on factors that are known to influence the pharmacokinetics of caffeine (e.g. smoking and oral contraceptive status). Often not even basic subject information (e.g. weight, sex, or age) are reported. These factors are essential in the analysis of the pharmacokinetics of any substance *in vivo* (Stader et al., 2019). In general-purpose pharmacokinetic data sets, concentration-time profiles are the fundamental and most valuable data type. Common practice however is not to report the raw measured data but only derived pharmacokinetic parameters or metabolic ratios. We strongly advocate the reporting of all data on an individual level while including detailed anonymized meta information alongside the concentration-time profiles. Access to the individual raw data would enable data integration with different data sets and the stratification of the data under various aspects. There are recent efforts in creating a standard resource for that matter (Grzegorzewski et al., 2020).

## CONFLICT OF INTEREST STATEMENT

The authors declare that the research was conducted in the absence of any commercial or financial relationships that could be construed as a potential conflict of interest.

## AUTHOR CONTRIBUTIONS

JG and MK designed the study, implemented and performed the analysis, curated the majority of caffeine studies and wrote the initial draft of the manuscript. FB and AK curated a subset of caffeine studies. All authors discussed the results. All authors contributed to and revised the manuscript critically.

## FUNDING

JG, FB, AK and JG were supported by the Federal Ministry of Education and Research (BMBF, Germany) within the research network Systems Medicine of the Liver (LiSyM, grant number 031L0054). MK, FB and AK were supported by the German Research Foundation (DFG) within the Research Unit Programme FOR 5151 “QuaLiPerF (Quantifying Liver Perfusion-Function Relationship in Complex Resection - A Systems Medicine Approach)” by grant number 436883643.

## DATA AVAILABILITY STATEMENT

The data sets analyzed for this study can be found in https://pk-db.com/.

## Permission to Reuse and Copyright

Figures, tables, and images will be published under a Creative Commons CC-BY licence and permission must be obtained for use of copyrighted material from other sources (including re-published/adapted/modified/partial figures and images from the internet). It is the responsibility of the authors to acquire the licenses, to follow any citation instructions requested by third-party rights holders, and cover any supplementary charges.

## ACKNOWLEDGMENTS

We acknowledge support by the BMBF-funded de.NBI Cloud within the German Network for Bioinformatics Infrastructure (de.NBI) (031A537B, 031A533A, 031A538A, 031A533B, 031A535A, 031A537C, 031A534A, 031A532B) for the PK-DB infrastructure.

## Notes

### Competing Interest Statement

The authors have declared no competing interest.

### Summary of Updates

Updated discussion section. Improved language. Additional data sets added. Additional feedback from coauthors included.

